# High-throughput multi-parallel enteropathogen quantification via nano-liter qPCR

**DOI:** 10.1101/746446

**Authors:** Jessica A. Grembi, Koshlan Mayer-Blackwell, Stephen P. Luby, Alfred M. Spormann

**Author notes:** Address correspondence to Jessica A. Grembi. Koshlan Mayer-Blackwell, *Fred Hutchinson Cancer Research Center, Vaccine and Infectious Disease Division, Seattle, WA, USA.

## Abstract

Quantitative molecular diagnostic methods, such as qPCR, can effectively detect pathogen-specific nucleic acid sequences. However, costs associated with multi-pathogen quantitative molecular diagnostics hinder their widespread use. Nano-liter qPCR (nL-qPCR) is a miniaturized tool for quantification of multiple targets in large numbers of samples based on assay parallelization on a single chip, with potentially significant cost-savings due to rapid throughput and reduced reagent volumes. We evaluated a suite of novel and published assays to detect 17 enteric pathogens using a commercially available nL-qPCR technology. Assay efficiencies ranged from 88-98% (mean 91%) and were reproducible across four operators at two separate facilities. When applied to complex fecal material, assays were sensitive and selective (99.8% of DNA amplified were genes from the target organism). Detection limits were 1-2 orders of magnitude higher for nL-qPCR than an existing enteric TaqMan Array Card (TAC), due to nanofluidic volumes. Compared to the TAC, nL-qPCR displayed 97% (95% CI 0.96, 0.98) negative percent agreement and 63% (95% CI 0.60, 0.66) overall positive percent agreement. Positive percent agreement was 90% for target concentrations above the nL-qPCR detection limits. nL-qPCR assays showed an underestimation bias of 0.34 log_10_ copies/gram of stool [IQR -0.41, -0.28] compared with the enteric TAC. Higher detection limits, inherent to nL-qPCR, do not hinder detection of clinically relevant pathogen concentrations. With 12 times higher throughput for a sixth of the per-sample cost of the enteric TAC, the nL-qPCR chip described here is a viable alternative for enteropathogen quantification for studies where other technologies are cost-prohibitive.

## Introduction

Quantitative molecular diagnostic methods, such as quantitative polymerase chain reaction (qPCR), can target nucleic acid gene sequences specific to known microbial pathogens. These methods have provided insights in the study of diarrheal disease beyond what can be gained using microbiological cell culture or immunoassays (1–4) and have been applied successfully in the field of pathogen detection for decades (5–9). Over time, molecular diagnostics were developed from single-gene qPCR assays to multiplex reactions (10–14) and to multi-assay, multi-sample arrays that can be operated in parallel on a single chip or card (15–19). Specifically in the field of enteric pathogen detection, a TaqMan Array Card (TAC) was developed by Liu and colleagues (15, 16) and subsequently used in several studies to estimate pathogen-attributable diarrhea burdens (3, 4, 20), as well as the impact of enteric pathogens on child growth (21–23) and vaccine uptake (24, 25). However, despite advances in the throughput of molecular detection of pathogens, costs associated with broad multi-target molecular assays still pose a barrier to their widespread use in epidemiological studies. For instance, the per-sample cost of the enteric TAC is $60-155, not including labor, capital equipment, nor DNA extraction reagents (15).

Compared with TaqMan qPCR arrays, higher-throughput microfluidic qPCR technologies hold potential to decrease per sample costs of multi-target diagnostics. In the case of nano-liter (nL) qPCR, precision robotic dispensing permits smaller reaction volumes, increases throughput, and reduces reagent volumes. While nL-qPCR technologies have been previously applied to pathogen detection, early efforts to develop nL-qPCR pathogen chips were limited by factors such as: (i) high-detection limits associated with small reaction volumes (6-33 nL), (ii) insufficient assay validation, and (iii) relatively low sample throughput per chip (12-48 samples) (26–28).

In more recent studies, a commercial nL-qPCR technology (SmartChip™ Real-Time PCR, TakaraBio Inc.) was used to design multi-target diagnostics to detect the presence of antibiotic resistance genes in urban wastewater treatment plant effluent, reclaimed water, and environmental samples (29–31) and to evaluate a suite of related dehalogenase genes in complex microbial communities (32). This technology uses 100 nL reaction volumes and allows for flexible configuration of a 5184-well chip that can analyze up to 384 samples (depending on the number of assays included). Using this platform, we developed a nL-qPCR chip with 54 assays (targeting 17 enteric pathogens) across 96 samples in duplicate. Here, we present comprehensive validation of the technology with laboratory standards as well as fecal samples from children in rural Bangladesh. The nL-qPCR enteropathogen chip permits high-throughput, rapid pathogen detection at significantly lower cost per-sample than other methods.

## Methods

### Assay design

We selected bacterial, protozoan, and helminthic enteropathogens identified as contributing to diarrheal disease in children across twelve countries (33, 34). We computationally designed and screened 175,000 candidate primer pairs to target 16 virulence genes using methods described previously (32). Briefly, amino acid sequences corresponding to all non-redundant members of each target gene’s protein family (Pfam v 27.0) were clustered based on percent pairwise identity using BLASTp all-vs-all search (35). We downloaded corresponding nucleotide sequences from NCBI, and DNA oligonucleotide primers were designed to target conserved DNA-level sequence motifs in sequence clusters containing the target virulence protein for each pathogen. A Python script directed the software primer3 (36) to develop thousands of candidate primer pairs for each target gene, which were then screened *in silico* against other non-target clusters within the same protein family. Up to eight assays per target gene were selected for laboratory screening. We included an additional 10 published assays (see Table 1) to assess the suitability for inclusion of previously validated assays optimized at similar PCR conditions (15, 37–41).

**Table 1.**
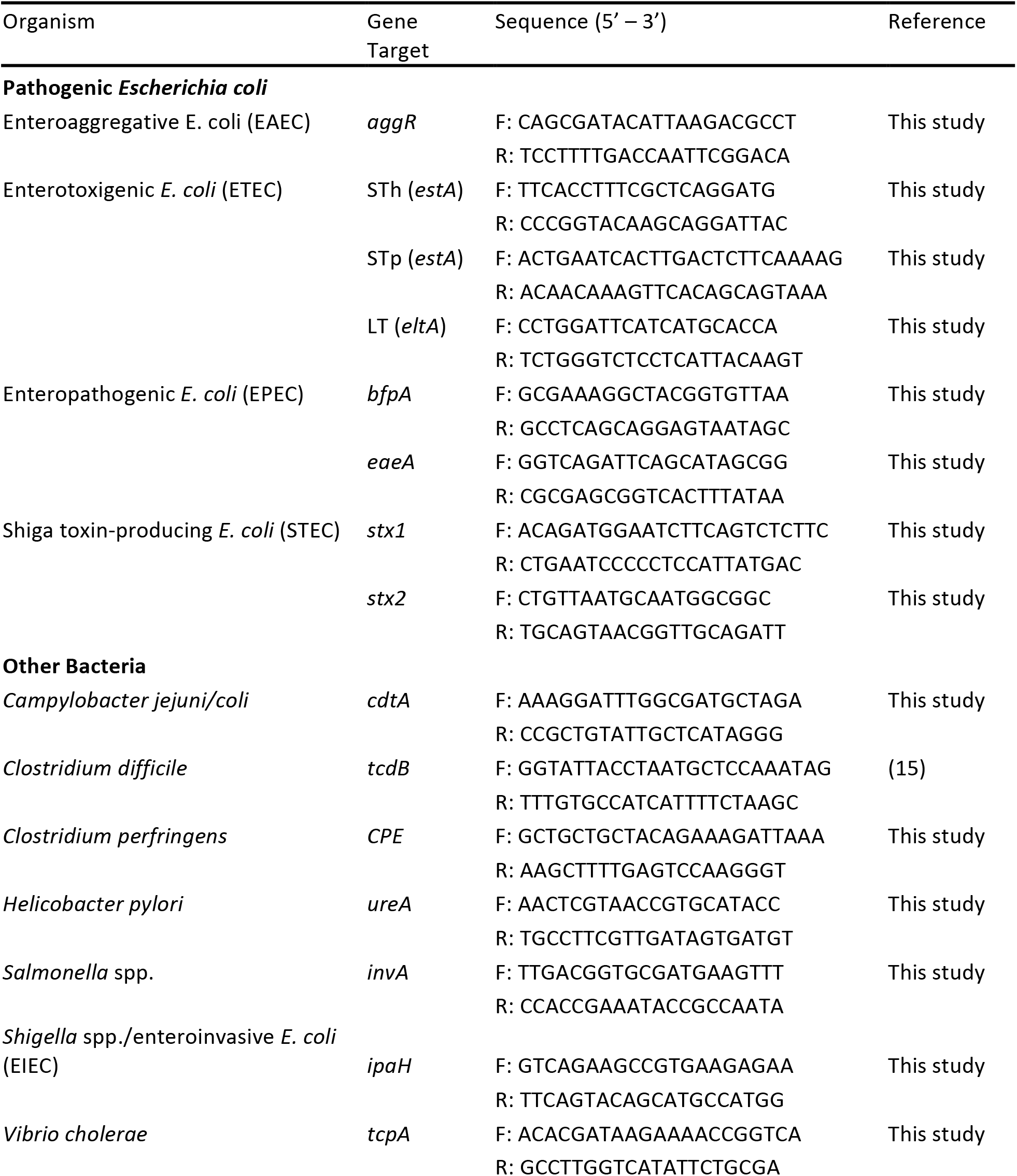
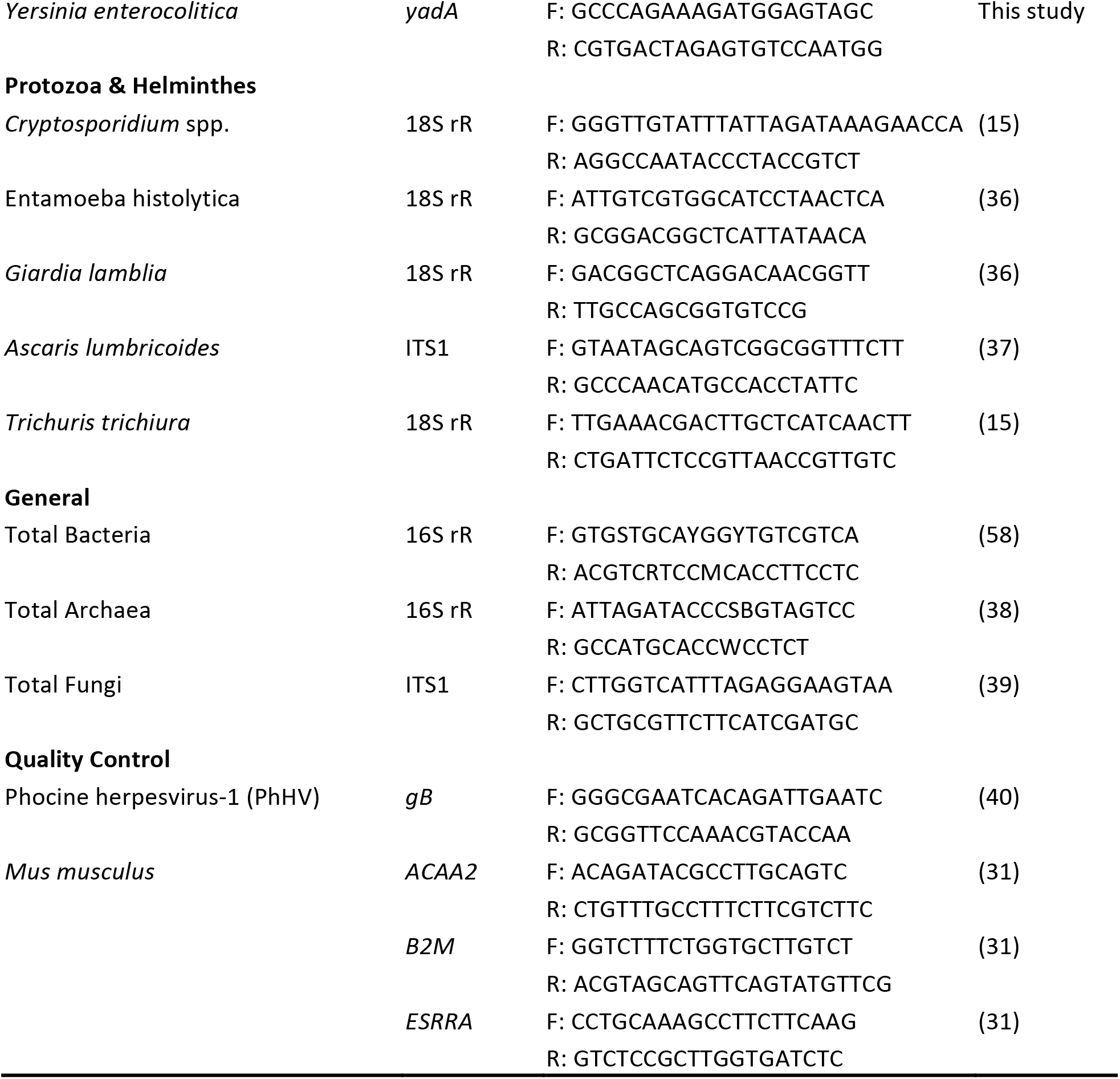
Assays included on the nL-qPCR pathogen chip. The chip contains each assay in duplicate, except for two of the Quality Control assays (*ACAA2* and *ESSRA*).

### nL-qPCR procedures and assay pre-screening

Assays were initially evaluated against 490bp synthesized linear DNA strands (Integrated DNA Technologies, Inc., Coralville, IA) for each target gene. Oligonucleotide primers (Integrated DNA Technologies, Inc., Coralville, IA) at a final concentration of 1 μM were added to LightCycler 480 SYBR Green I Master Mix (Roche Applied Sciences, Indianapolis, IN) and were robotically dispensed onto nL-qPCR chips using TakaraBio’s SmartChip™ platform. In a separate plate, samples were added to additional master mix and robotically dispensed onto chips. Duplicate chips were run, using the standard TakaraBio protocol: 95°C for 3 min, then 40 cycles of (95°C for 60s, 60°C for 70s). We excluded assays that reproducibly displayed fluorescence in the negative control (PCR grade water) prior to cycle 28, failed to amplify standards at 100 copies per well, or had PCR efficiencies less than 85%. Final assays were selected based on optimal performance characteristics as described below. Each chip contained a minimum of two negative (no-template) controls for each assay.

### Analytical performance characteristics

Analytical performance was evaluated in accordance with the Minimum Information for Publication of Quantitative Real-Time PCR Experiments (MIQE) guidelines (42). Assay efficiencies were evaluated with a pool of synthetic DNA standards, described above. Standards were 10-fold serially diluted (10 to 10^6^ copies/reaction). Standard curves were run on a minimum of 15 chips over two instruments at separate facilities (Fremont, CA and East Lansing, MI) and with two different operators at each location. Efficiencies were calculated according to Rutledge and Côté (43); mean efficiency over all runs is reported along with coefficient of variation (CV). Limit of detection (LOD) was determined with pooled synthetic DNA standards spiked into extracted DNA from 10 fecal samples to a final concentration of 10, 100, and 1000 copies/reaction; then each sample was run in duplicate on two separate chips. The mean cycle quantification (C_q_) value (i.e. the cycle at which sufficient copies of target DNA have been made to produce a fluorescent signal detectable by the instrument) was calculated for duplicate assays on a single chip, and all results under the C_q_ cutoff of 30 were determined positive. A total of 20 positive samples (10 samples x 2 chips) per target at each concentration were assayed and LOD is defined as the lowest concentration which 95% were positively detected (i.e. where 19 of the 20 were detected).

Inter-assay precision (reproducibility) was assessed across the standard curves used for efficiency calculations measured over 15-20 chips, using different lots of master mix, different batches of oligonucleotide primers, and 4 different operators at 2 separate facilities. We report the mean CV on calculated copy numbers over all points on the standard curve as well as the range. Intra-assay precision (repeatability) was measured within-chip and between chips. Within-chip precision was evaluated in three samples in which extracted DNA from fecal samples was mixed with positive controls at high (10^5^ copies/reaction) and low (100 copies/reaction) concentrations and assayed 20 times on a single chip: we report the CV of calculated copy number across the 20 replicates. Between-chip precision was evaluated in 223 fecal samples collected from a cohort of Bangladeshi children that tested positive for at least one pathogen (C_q_ < 30) plus 18 additional fecal samples into which we added positive controls. Replicates for each sample were run on two chips and the CV of calculated template copies was determined across all four replicates. We report mean CV of calculated template copies over all samples, as well as the number of unique samples included in the calculation of the mean.

Sensitivity and specificity were evaluated using four pools of DNA standards, spiked into extracted DNA from 40 pathogen-free fecal samples. For each pathogen 10 samples contained the target at low concentration (100 copies/reaction), 10 samples at medium concentration (10x the LOD) and 10 samples at high concentration (100x the LOD); an additional 10 samples had no target. Sensitivity and specificity were determined based on positive or negative detection in these 40 samples. In order to further verify assay specificity, we sequenced PCR amplicons obtained from running 96 child fecal samples on the nL-qPCR chip. The Seq-Ready™ TE MultiSample FLEX protocol, PCR clean-up, and DNA quantification prior to sequencing were done in accordance with TakaraBio’s standard procedures, as described previously (Atshemyan, 2017; Firtina, 2017). The resulting paired-end Illumina MiSeq reads were quality filtered and only sequences that were the expected target gene amplicon length (+/− 3 bp) were maintained. We verified the intended target (organism and gene) by conducting a nucleotide BLAST search (35) on each unique sequence. We retained the top hit(s), defined as the highest sequence identity with the lowest E value.

### Sample collection

To test the performance of nL-qPCR chip and against the performance of enteric TAC in epidemiology-relevant samples, we utilized 254 fecal samples from children in rural Bangladesh. Children were between 10 and 18 months old and enrolled in the WASH Benefits randomized controlled trail (44–47). Split samples from these children had been previously assayed for pathogens with the TAC technology (manuscript *in preparation*). Samples were collected by the child’s caregiver into a sterile collection container and placed on cold chain within 165 [IQR 79, 791] minutes, transported to the laboratory and held at −80°C prior to analysis. DNA was extracted according to previously published protocols (16) in the Parasitology lab at the International Centre for Diarrhoeal Disease Research, Bangladesh (icddr,b) and separated into two aliquots: one aliquot was subjected to TAC analysis at icddr,b and the other was shipped on dry ice to Stanford University. Samples were collected after obtaining written, informed consent from the child’s primary caregiver and with approval from human subjects committees at icddr,b (PR-11063), University of California, Berkeley (2011-09-3652), and Stanford University (25863).

### Statistical analyses

Data analysis was performed in R statistical software (v3.5.2) and analysis files are available as supplemental files. CV, the standard deviation of replicates divided by the mean, was used to evaluate precision in accordance with the MIQE guidelines (42). Specifically, CV of calculated copy number, and not C_q_ value, is reported per Schmittgen and Livak (48) and Hellemans et al. (49). Within-chip, between-chip, and between instrument/operator variances were compared with a pairwise Wilcoxon rank sum test, using the Benjamini-Hochberg procedure to account for multiple comparisons (50). Sensitivity and specificity were calculated using the **epi.test** function from the **epiR** package (51). Positive percent agreement and negative percent agreement were calculated in the same manner and are reported with this alternative nomenclature as recommended when no known reference standard is used (52). Exact binomial 95% confidence limits on sensitivity and specificity were calculated according to Collett (53). Unweighted Cohen’s Kappa was calculated using the **epi.kappa** function with confidence intervals calculated according to Rothman (54). Bias in calculated log_10_ copy numbers per gram of stool (corrected for extraction and PCR efficiency by normalizing to the positive control PhHV spike-in) was evaluated according to Bland and Altman (55) using the **blandr::blandr.statistics** function to estimate bias (56); 95% confidence intervals determined per Bland and Altman (57).

### Data Availability and Cost Estimates

Nucleotide sequences obtained from Bangladeshi child fecal samples for the specificity analysis are deposited in the Sequence Read Archive under BioProject # SUB5519617. Cost estimates for both nL-qPCR and TAC qPCR technologies can be found in Table S1.

## Results

### Analytical performance

The mean efficiency for each assay, based on the evaluation of standard curves run on 15-20 chips, ranged from 88-98% (mean 91%) with a coefficient of variation of 6.3% [IQR 5.3, 7.3](Table 2). The linearity over all assays on all chips was 0.990 [IQR 0.987, 0.992] and detection limits were between 10-100 copies/100nL reaction, which corresponds to 8×10^5^-8×10^6^ copies/g of stool (Table 2). Within-chip repeatability was assessed in ten replicates on a single chip: synthetic DNA in high (10^5^ copies/reaction) and low (10^2^ copies/reaction) concentrations was spiked into DNA extracted from fecal samples. The high concentration displayed a coefficient of variation in calculated copy number of 15% [IQR 8 – 25]); the low concentration had variability of 27% [IQR 18 – 36] (Figure S1).

**Table 2.** Analytical performance of the nL-qPCR pathogen chip.

| Organism (gene target) | Efficiency<br>% (CV<sup>a</sup>) | LOD<sup>b</sup><br>(copies/reaction) | Reproducibility<br>CV above, at LOD<sup>c</sup> | Repeatability<br>CV (n)<sup>d</sup> | Sensitivity | Specificity |
| --- | --- | --- | --- | --- | --- | --- |
| **Pathogenic E. coli** |  |  |  |  |  |  |
| EAEC (aggR) | 95 (0.16) | 8e+05 (10) | 26, 115 | 19 (75) | 100 | 93 |
| ST-ETEC (STh) | 90 (0.07) | 8e+06 (100) | 22, 51 | 44 (8) | 100 | 100 |
| ST-ETEC (STp) | 92 (0.08) | 8e+06 (100) | 28, 39 | 28 (13) | 100 | 100 |
| LT-ETEC (eltA) | 91 (0.08) | 8e+06 (100) | 24, 49 | 18 (25) | 100 | 100 |
| EPEC (bfpA) | 89 (0.05) | 8e+06 (100) | 26, 43 | 32 (13) | 100 | 100 |
| EPEC (eaeA) | 90 (0.05) | 8e+05 (10) | 23, 55 | 33 (59) | 100 | 93 |
| STEC (stx1) | 89 (0.08) | 8e+06 (100) | 18, 47 | 46 (6) | 100 | 100 |
| STEC (stx2) | 92 (0.08) | 8e+06 (100) | 22, 57 | 62 (16) | 100 | 100 |
| **Other Bacteria** |  |  |  |  |  |  |
| Campylobacter jejuni/coli (cdtA) | 92 (0.07) | 8e+06 (100) | 32, 52 | 29 (35) | 100 | 100 |
| Clostridium difficile (tcdB) | 90 (0.05) | 8e+06 (100) | 20, 34 | 25 (8) | 100 | 100 |
| Clostridium perfringens (CPE) | 88 (0.06) | 8e+05 (10) | 22, 52 | 24 (2) | 98 | 100 |
| Helicobacter pylori (ureA) | 91 (0.07) | 8e+06 (100) | 28, 92 | 42 (2) | 100 | 100 |
| Salmonella enterica (invA) | 91 (0.07) | 8e+05 (10) | 21, 53 | 68 (3) | 100 | 100 |
| Shigella/EIEC (ipaH) | 90 (0.07) | 8e+06 (100) | 20, 29 | 25 (15) | 100 | 100 |
| Vibrio cholerae (tcpA) | 89 (0.06) | 8e+06 (100) | 20, 44 | 41 (1) | 100 | 100 |
| Yersinia enterocolitica (yadA) | 92 (0.07) | 8e+06 (100) | 26, 61 | 64 (2) | 100 | 100 |
| **Protozoa & Helminthes** |  |  |  |  |  |  |
| Cryptosporidium (18S) | 88 (0.07) | 8e+06 (100) | 36, 103 | 76 (4) | 100 | 100 |
| Entamoeba histolytica (18S) | 90 (0.1) | 8e+06 (100) | 28, 46 | 54 (2) | 100 | 100 |
| Giardia (18S) | 90 (0.07) | 8e+06 (100) | 21, 31 | 33 (17) | 100 | 90 |
| Ascaris lumbricoides (ITS1) | 89 (0.05) | 8e+05 (10) | 20, 53 | 16 (2) | 100 | 100 |
| Trichuris trichiura (18S) | 91 (0.07) | 8e+05 (10) | 17, 58 | 28 (2) | 100 | 100 |
| **General** |  |  |  |  |  |  |
| Total Archaea (16S) | 90 (0.06) | 8e+06 (100) | 44, 75 | 70 (2) | 100 | 93 |
| Total Bacteria (16S) | 98 (0.09) |  | 31, 319 |  |  |  |
| Total Fungi (ITS1) | 91 (0.1) | 8e+06 (100) | 43, 54 | 58 (34) | 100 | 87 |
| **QC** |  |  |  |  |  |  |
| PhHV (gB) | 92 (0.07) | 8e+06 (100) | 22, 66 | 52 (92) | 100 | 100 |

<sup>a</sup> Coefficient of variation (CV) of efficiency calculated across 15-20 chips
<sup>b</sup> Minimum no. of gene copies per g of stool (minimum no. gene copies per 100 nL reaction)
<sup>c</sup> CV on calculated copy number for all points along the standard curve measured over 15-20 chips, shown separately for concentrations above the LOD and at the LOD
<sup>d</sup> CV in calculated copy number across 4 replicates as measured in n positive samples, mean CV is reported

C_q_ values across replicate chips were highly repeatable for synthetic DNA standards (R^2^ = 0.989, Figure 1a), for synthetic DNA in a complex stool DNA matrix (R^2^ = 0.984, Figure 1b) and for DNA extracted from fecal samples collected from children in Bangladesh (R^2^ = 0.935, Figure 1c). Fecal samples displayed a median difference in C_q_ values of 0.39 [IQR 0.15 – 0.81](Figure 1c) across all assays, which corresponds to a coefficient of variation on calculated gene copy number of 28% [IQR 16 – 50] (Table 2, Repeatability). The highest variability was again seen at the lowest concentrations (Figure S2).

**Figure 1.**
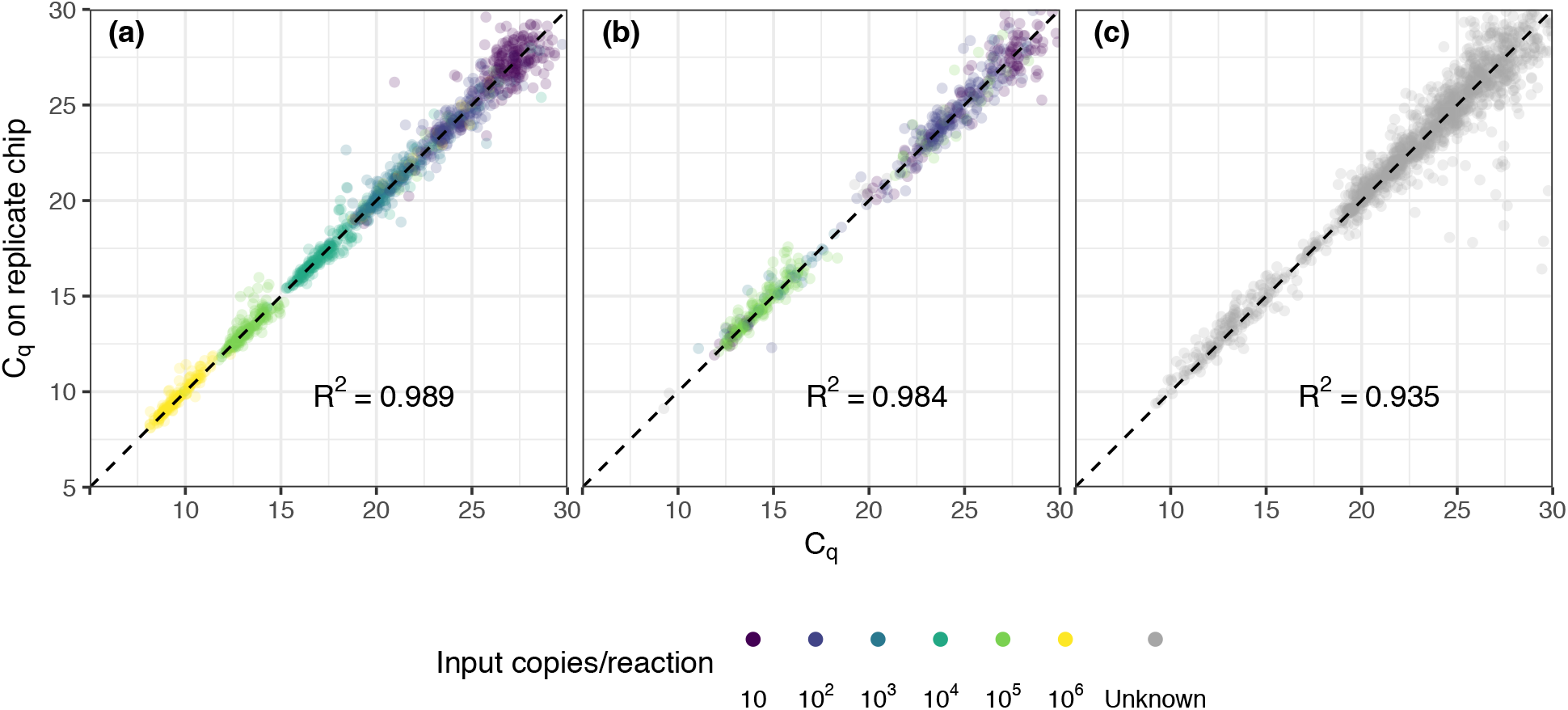
Assay precision across replicate chips for (a) synthetic DNA standards across a 6-fold dilution series, (b) synthetic DNA standards spiked into child fecal samples (n = 60), and (c) child fecal samples (n = 254). Each point represents the replicated results for a single sample-assay reaction run in a specific location on the chip. Points shown with color indicate results from amplified DNA standards with defined input copy number (10-10^6^); grey points indicate results from fecal samples with unknown input copy number, absent synthetic standards.

Assays were reproducible across two instruments and four operators, again with an inverse relationship noted between variance and concentration (Table S2). At concentrations one or more orders of magnitude above the detection limit, coefficient of variation on calculated copy number ranged from 17 – 44% (Table 2). Coefficient of variation at the limit of detection ranged from 29% to 115% for pathogen virulence and marker genes, the highest of which was analogous to 17 ± 20 copies detected. The highest variance (319%) observed was for the total bacterial (16S rRNA) assay at the detection limit of 10 copies/reaction. Between-chip variance was similar to variance across two instruments and four operators (p = 0.99) but both were significantly higher than within-chip variance (p < 0.0001, pairwise Wilcoxon rank sum test). Coefficients of variation of the magnitudes observed are not biologically relevant when analyzing pathogen quantities on the log_10_ scale, as is the normal procedure.

Analytical sensitivity ranged from 98-100% and specificity from 90-100% (Table 2) among 60 samples containing combinations of synthetic nucleic acid spiked into DNA extracted from 15 different individuals and assayed in duplicate. To further ensure the specificity of the assays, we sequenced amplicons from 96 fecal samples collected from children in Bangladesh that tested positive for at least one pathogen target. We obtained 1.7M (26,747 unique) sequences with 330 [IQR 142, 1171] unique sequences per assay. Amplicon sequencing showed that the assays were specific. The intended gene target was correctly identified in the top hit(s) (defined as highest identity and lowest E value) for 99.8% of unique sequences. Most (99.7%) of the BLASTn searches returned a database top hit with ≥97% sequence identity. The *Ascaris lumbricoides* assay had highest number of off-target hits: 7/130 of the unique sequences were identified as the same target gene in a closely related species, *Ascaris ovis*.

### Clinical performance

We analyzed 254 fecal samples collected from children in Bangladesh on both the nL-qPCR chip and the TAC to compare performance. Overall percent agreement was 90% for the >4500 reactions and negative percent agreement was 97% (95% CI 0.96, 0.98)(Cohen’s Kappa = 0.66 (95% CI 0.63 – 0.69)). Positive percent agreement was highly dependent on concentration of the target gene. At concentrations above nL-qPCR detection limits (>10^7^ copies/g stool) positive percent agreement was 90%; this dropped to 62% for concentrations near the nL-qPCR detection limits (10^5^-10^7^ copies/g stool) and fell to 8% for concentrations below 10^5^ copies/g stool. In instances where both methods detected the presence of target genes, nL-qPCR assays displayed a median underestimation bias of −0.34log_10_ copies [IQR -0.41, -0.28] (see Table 3 and Figure S3 for individual assay statistics).

**Table 3.**
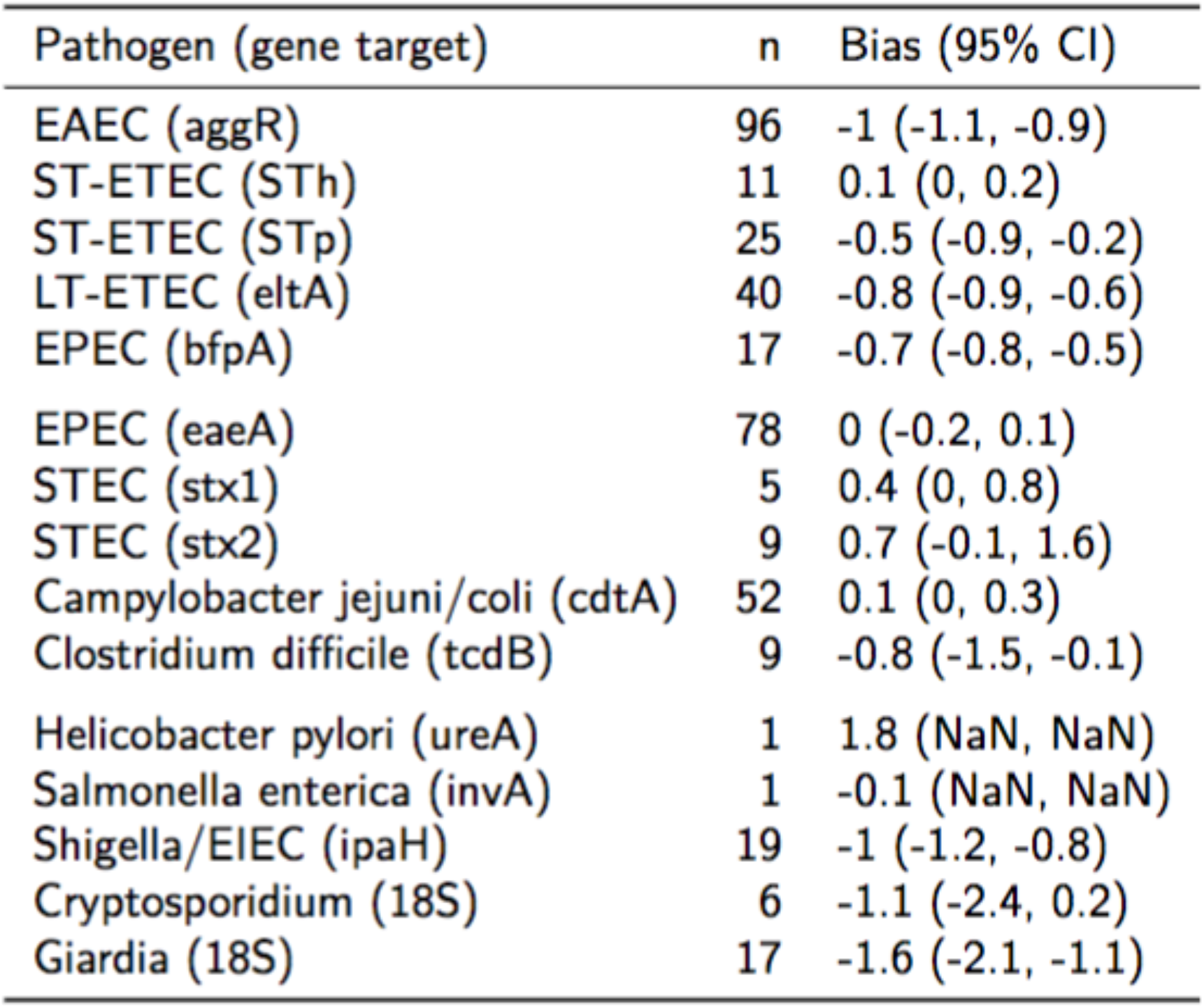
Bias estimates by assay on calculated log_10_ copy number per gram of stool for nL-qPCR compared to TAC. The table includes 529 reactions that were concordant for detection on both platforms.

| Pathogen (gene target) | n | Bias (95% CI) |
| --- | --- | --- |
| EAEC (aggR) | 96 | -1 (-1.1, -0.9) |
| ST-ETEC (STh) | 11 | 0.1 (0, 0.2) |
| ST-ETEC (STp) | 25 | -0.5 (-0.9, -0.2) |
| LT-ETEC (eltA) | 40 | -0.8 (-0.9, -0.6) |
| EPEC (bfpA) | 17 | -0.7 (-0.8, -0.5) |
| EPEC (eaeA) | 78 | 0 (-0.2, 0.1) |
| STEC (stx1) | 5 | 0.4 (0, 0.8) |
| STEC (stx2) | 9 | 0.7 (-0.1, 1.6) |
| Campylobacter jejuni/coli (cdtA) | 52 | 0.1 (0, 0.3) |
| Clostridium difficile (tcdB) | 9 | -0.8 (-1.5, -0.1) |
| Helicobacter pylori (ureA) | 1 | 1.8 (NaN, NaN) |
| Salmonella enterica (invA) | 1 | -0.1 (NaN, NaN) |
| Shigella/EIEC (ipaH) | 19 | -1 (-1.2, -0.8) |
| Cryptosporidium (18S) | 6 | -1.1 (-2.4, 0.2) |
| Giardia (18S) | 17 | -1.6 (-2.1, -1.1) |

Reactions detected by TAC but not by nL-qPCR were typically below nL-qPCR detection limits, with TAC C_q_ values approximately 30.9 [IQR 28.6, 33.0] (Figure 2a, black points). The higher detection limits for nL-qPCR assays did not interfere with detection of diarrhea-causing pathogen concentrations, with the exception of the *V. cholerae* assay which had an etiologic cutoff that was below the nL-qPCR detection limit. The etiologic cutoff (shown as red lines in Figure 2a) indicates the TAC C_q_ value below which children were highly likely to have diarrhea, i.e. the value at which the odds ratio for diarrhea cases compared to controls was greater than 2 (Liu, 2016, Platts-Mills, 2018). nL-qPCR assays detected all but 8 of the 40 reactions in which TAC assays detected a sample below the etiologic C_q_ cutoff value (3 of which were for *V. cholerae*), and typically detected samples well above the cutoff for most assays (Figure 2a). Reactions positive by nL-qPCR but not TAC were also at low concentrations (Figure 2b) and could have been the result of less stringent amplification without the use of probe-based dyes with nL-qPCR.

**Figure 2.**
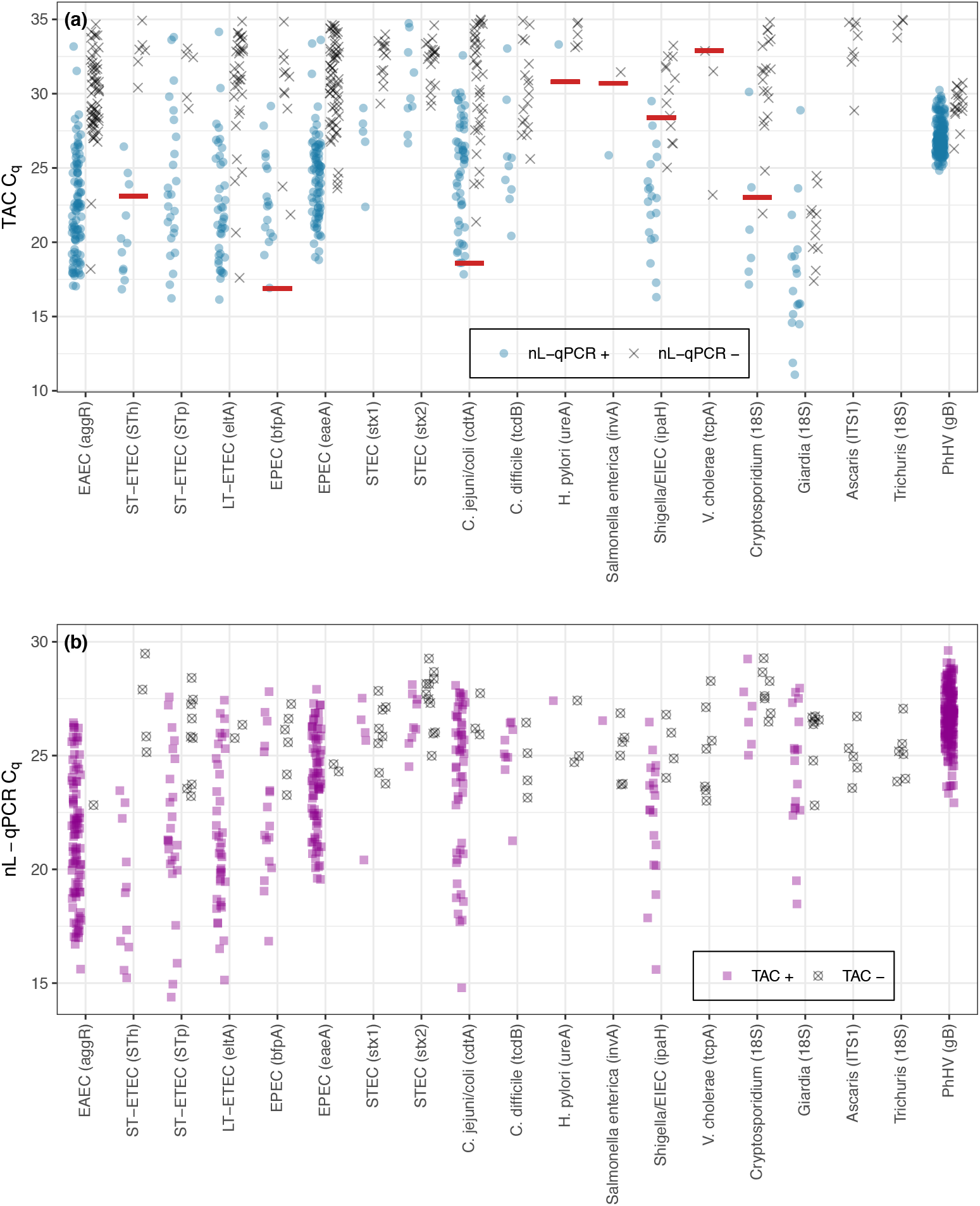
Comparison of nL-qPCR and TAC assays across 254 fecal samples. Samples detected by (a) TAC or (b) nL-qPCR are shown with their respective C_q_ values. Blue (top) and purple (bottom) points represent positive detections found in both nL-qPCR and TAC tests. Gray points (top) represent targets undetected by nL-qPCR but detected by TAC, and vise versa (bottom). Red lines represent pathogen TAC etiologic cutoff C_q_ values (etiologic cutoffs have not been established for nL-qPCR assays).

Contamination may cause false-positive qPCR results, and can occur due to cross-contamination between samples or as a result of free ambient DNA in the laboratory environment. Sample cross-contamination occurred rarely with nL-qPCR; amplification of pathogen virulence or marker genes in no-template controls occurred in <3% of the 4288 no-template control sample reactions. Moreover, these amplifications resulted in calculated copy numbers near or below the established limit of detection (median 35 [IQR 28 – 42] calculated copies). Ambient laboratory contamination was detected more frequently. Amplification of bacterial 16S rRNA occurred in 46% of no-template controls, and was highly dependent on operator. In all cases, contamination was near the detection limit with 11 [IQR 7, 25] calculated copies of bacterial 16S rRNA detected. Cross-contamination, although possible, occurs rarely and only in low concentration, thereby indicating a low likelihood of false positive results.

## Discussion

The nL-qPCR chip evaluated here provides satisfactory analytical performance for simultaneous analysis of 96 samples against a suite of 17 enteric pathogens for a cost of <$10/sample. The high-throughput nature of the nL-qPCR chip is particularly advantageous when large numbers of samples need to be processed in a timely manner, such as in population-based studies. Above certain thresholds, we found analytical performance to be comparable to an enteric TAC widely used for investigations of diarrheal disease in diverse global populations, for roughly a quarter of the per-sample cost ($60; (15)).

The primary difference in performance we observed was that a majority of the nL-qPCR assays had detection limits 1-2 orders of magnitude higher than TAC. This was due to reactions that utilize 16 times less sample volume (0.0125 μL compared to 0.2-0.4 μL for TAC; (15,personal communication with J. Liu, 2019). Furthermore, among 254 fecal samples from Bangladeshi children 14 months old, most nL-qPCR assays displayed an underestimation bias (i.e. returned a lower estimated number of copies per gram of stool) compared to the enteric TAC. However, these differences do not appear to be limitations in terms of ability to distinguish pathogen loads relevant for diarrheal disease for pathogens with etiological cutoffs established, with the potential exception of infection with *Vibrio cholerae*. Importantly, the TAC and nL-qPCR assays for *V. cholerae* target different virulence genes: hemolysin (*hlyA*) for TAC and toxin-coregulated pilus (*tcpA*) for nL-qPCR. The etiologic cutoffs were established for *hlyA*, which is commonly detected in environmental *V. cholerae* strains that lack both the *tcpA* and cholera toxin genes (58). Thus, discordant detection between the technologies might not represent differences in performance, but rather differential presence of these virulence genes within *V. cholerae* strains. Given that studies have shown low concentrations of *V. cholerae hlyA* gene are observed in feces coincident with diarrheal symptoms in children (3, 4), this might be a superior gene target for *V. cholerae* in pathogen panels. Additional studies should verify the optimal gene target for diarrhea-causing *V. cholerae* species.

In studies where quantitation is required at lower concentrations than were achieved in this study, pre-amplification can be performed as described by Ishii et. al. (28). In addition, pre-printing primers directly onto chips, similar to the TAC spotting procedure, can reduce detection limits by nearly 50%. However, a major advantage of the nL-qPCR SmartChip™ is the flexibility of the platform. Therefore, if a research team does opt to pre-print primers onto chips, we suggest also maintaining a stock of unprinted chips on-hand. The current configuration of the chip was designed with large-scale epidemiology studies in mind, thus increased throughput was prioritized over the inclusion of a higher number of assays. However, researchers wishing to focus on a smaller set of targets can evaluate more samples per plate (further reducing per-sample costs), or the number of samples can be reduced to accommodate an increased number of assay targets. In large-scale studies, replicating analysis for questionable samples is often necessary (e.g. when replicates give discordant results). Unprinted chips allow for an operator to run a limited suite of sample/assay pairs that need to be reanalyzed: for example, 384 samples with questionable results in the initial run from a large study could be analyzed against a minimal suite of 12 assays on a specially designed chip at the end of the study. This facilitates the resolution of discordant results and minimizes missing values in the final dataset, which will maximize statistical power in the analysis stage.

Unprinted nL-qPCR chips also allow end-users to substitute assays from the ones that we publish here, with appropriate assay validation. This evaluation included 10 pre-published assays that operate at similar PCR conditions, and found they performed well in nL format, suggesting end users have flexibility in re-designing the chip. We further show that seven primer pairs previously validated using TaqMan with probe-based dyes had excellent specificity among 96 fecal samples when utilized with SYBR Green intercalating dye instead. These results suggest the additional reagent costs associated with probes is not necessary to achieve high specificity and is consistent with other findings that have reported equal or superior specificity with SYBR Green compared to TaqMan chemistry (59, 60).

Quantifying nucleic acid targets for large numbers of samples is costly, regardless of the platform used, and recommended best practices are sometimes sacrificed in the face of limited budgets. For example, technical replicates are generally encouraged to facilitate identification of outlier or spurious results, particularly on chip- or card-style platforms, and increase the likelihood of detection near the detection limit where analytical precision is the lowest (15, 61, 62). The nL-qPCR pathogen chip is configured to provide duplicate results for the 24 pathogen-specific virulence and marker genes. This was deliberate as it is impossible to determine *a priori* if a sample will be near the detection limit, particularly in the case of fecal samples where the presence of PCR inhibitors is likely (63, 64). Early versions of the enteric TAC included replicates (15), but those have been replaced by additional pathogen targets in latter versions currently in use for large-scale studies (3, 4). Due to the flexibility in configuration of the nL-qPCR, up to 13 additional pathogen targets could be added without sacrificing duplicate assays, and throughput would still be 8-9 times higher and cost 50% less than the enteric TAC. The lower per-sample cost reduces the temptation to compromise best practices in the face of budgetary constraints.

The nL-qPCR platform has important limitations. First, due to the open chip technology, there is higher likelihood for contamination if not used in a controlled laboratory with minimal ambient contamination and meticulous operators. nL-qPCR does not appear to be well-suited for absolute quantification of total bacteria due to the fact that general bacterial contamination (via 16S rRNA) was detected in almost half of the no-template control samples, albeit at concentrations near the detection limit. To ensure potential low-concentration contamination is identified, we strongly recommend incorporation of replicates when using this technology or more stringent C_q_ filtering (e.g. C_q_ 28 or lower). Another limitation of the current configuration is the omission of viral enteric pathogen targets. The primary aim for this study was to validate the nL-qPCR technology for bacterial and parasitic targets, and we expect that future iterations of the chip will include viral targets, which could be combined with a reverse-transcriptase protocol for the study of RNA as well as DNA viruses.

In conclusion, we found the nL-qPCR pathogen chip to be an acceptable alternative to other methods; particularly for studies with large numbers of samples as savings in both cost and time will be amplified at scale.

## Acknowledgements

We would like to thank Joerg Deutzmann for molecular insights; and Albert Mueller and Wenyu Gu for critical comments on early versions of the manuscript. We wish to acknowledge Max Sanchez and Masy Leung (TakaraBio) as well as Jeff Landgraf and Christi Harris (Research Technology Support Facility, Michigan State University) for performing microfluidic reactions. We are grateful to the Stanford Health Care Clinical Microbiology Laboratory, and the laboratories of Manuel Amieva, Alexandria Boehm, Rashidul Haque, Justin Sonnenburg, David Stevens, and Steven Williams for providing pure culture DNA for select pathogens and/or positive control samples.

Funding for this work was provided by a grant from the National Institutes of Health (C-IDEA 1RC4TW00878101 to JAG and AMS) and a grant from the Bill and Melinda Gates Foundation (OPP1161946 to SPL). AMS was supported by a National Science Foundation (NSF) grant (C-DEBI STC). JAG was supported by a NSF Graduate Research Fellowship and a Stanford Interdisciplinary Graduate Fellowship. Funders had no role in the design or execution of the study, interpretation of results, or decision to publish.

---

**Figure S1.**
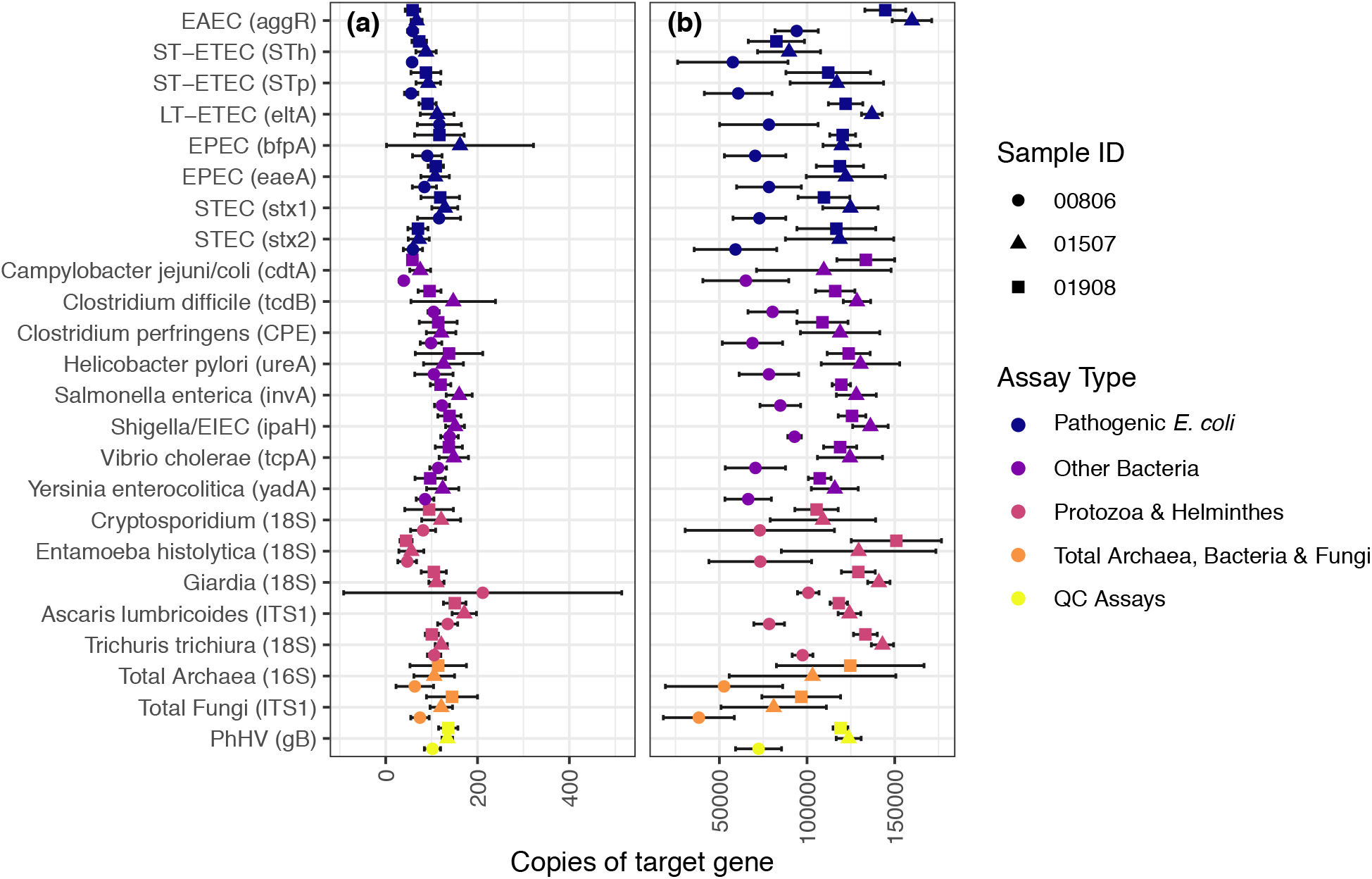
Within-chip intra-assay precision among samples containing a mixture of synthetic DNA standards and extracted DNA from 3 separate fecal samples. Each sample was assayed 10 times on a single chip. Synthetic DNA was added at either (a) 100 or (b) 10^5^ copies/reaction. Points represent the mean value across the 10 replicates and error bars show the standard deviation. The total bacteria results are not shown as fecal samples contained an unknown quantity of bacterial 16S rRNA which confounds the interpretation.

**Figure S2.**
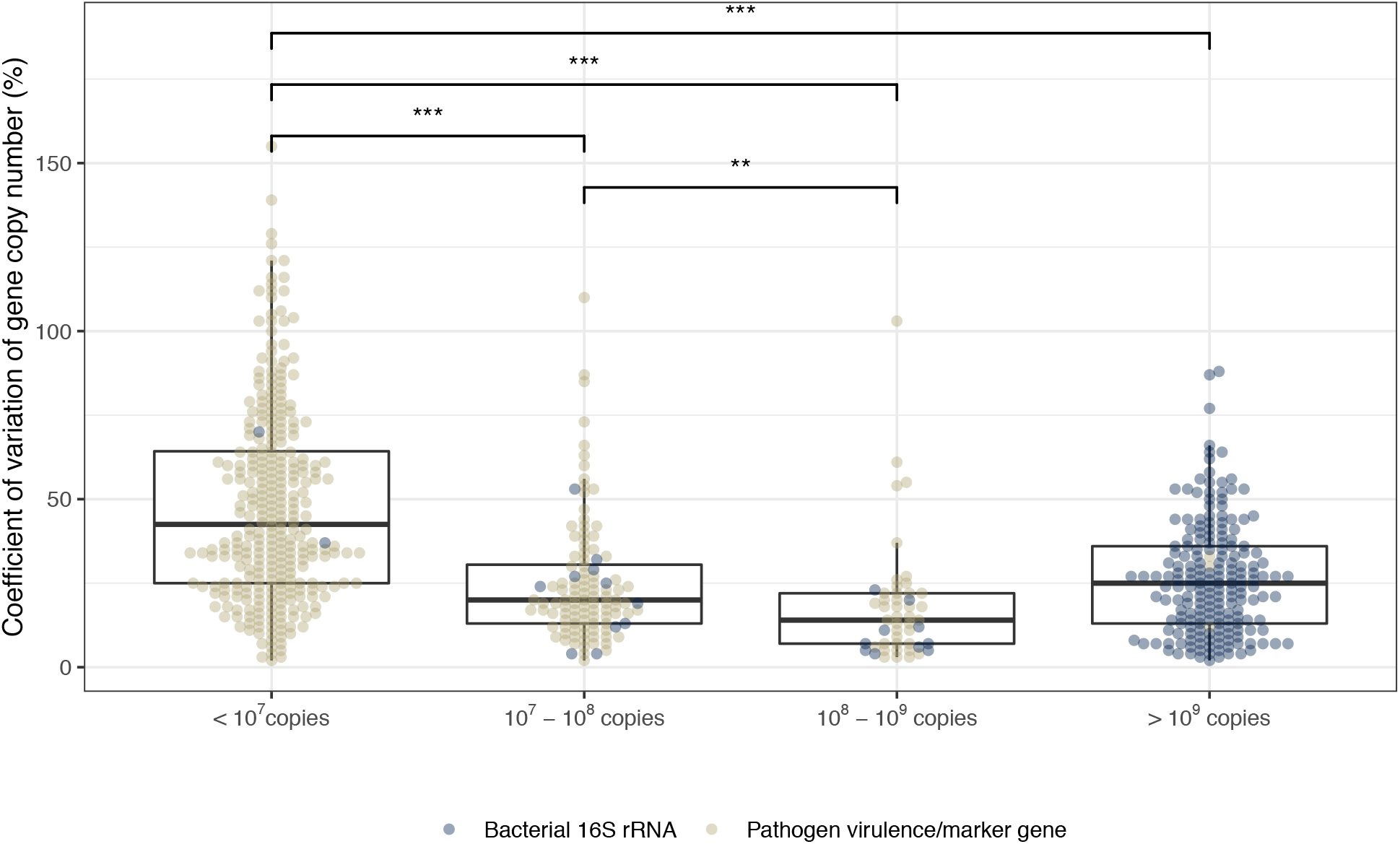
Calculated copy numbers have the highest variation at the lowest concentrations. Boxplots show median value and inner quartile ranges, points represent the result for a single sample-assay reaction run in quadruplicate (twice each on two chips), colors indicate if the assay targeted the general bacterial 16S rRNA gene or a specific pathogen virulence/marker gene. *** p < 0.001, ** p < 0.01 as determined by Wilcoxon rank sum test.

**Figure S3.**
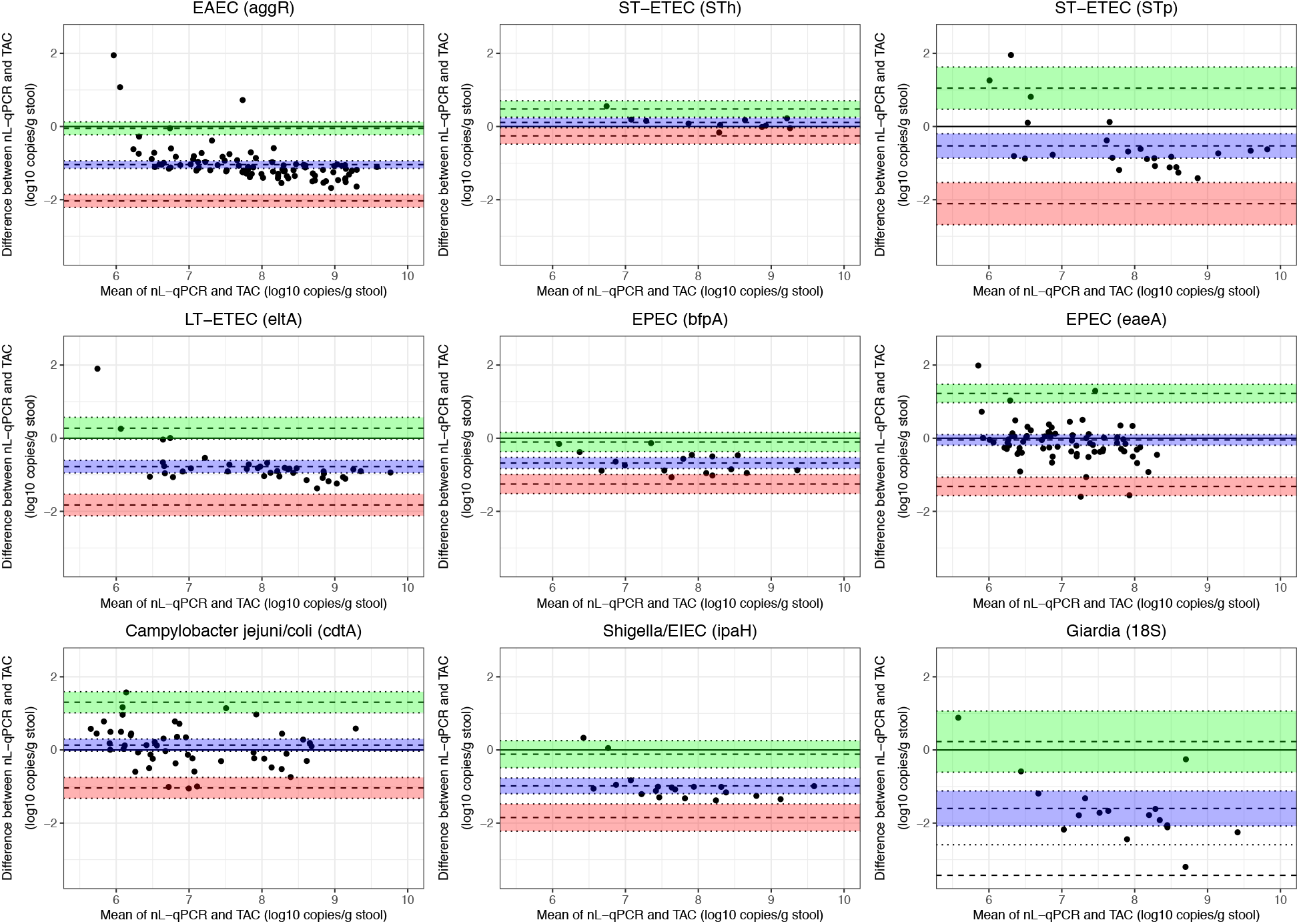
Bland Altman plots for assays with at least 10 detections in fecal samples in both nL-qPCR and TAC. The blue shaded area represents the mean bias and its 95% confidence interval. The upper (green) and lower (red) limits of agreement and their corresponding 95% confidence intervals are also shown.

**Table S1.** Cost comparison of consumable supplies for nL-qPCR enteric pathogen chip and enteric TaqMan Array Card (TAC). Nonconsumable costs include a specialized thermocycler instrument (QuantStudio for TAC; SmartChip Cycler for nl-qPCR) and additional instruments for sample handling (centrifuge with TAC-supported bucket adapters for TAC; SmartChip MultiSample Nano Dispenser robotic fluid handling system for nL-qPCR). Cost estimates obtained from Stanford University purchasing system in August 2019.

| Characteristic | nL-qPCR pathogen chip | Enteric TAC |
| --- | --- | --- |
| No. reaction wells | 5184 | 384 |
| No. samples (per chip or card) | 96 | 8 |
| Cost (per chip or card) | \$475 | \$600 |
| Mastermix<sup>a</sup> cost (per sample) | \$1.30 | \$78 |
| Primer<sup>b</sup> cost (per sample) | \$0.43 | NA (spotted on TAC) |
| Consumables-tips, tubes, 96-well plates (per sample) | \$3.00 | \$1.50 |
| **TOTAL (per sample)** | **\$9.68** | **\$155** |

<sup>a</sup> AgPath-ID One-Step RT-PCR Kit (1000 reactions; \$1600) for TAC; Roche LightCycler 480 SYBR Green I (103680 reactions; \$2500) for nL-qPCR
<sup>b</sup> IDT primer plate with mixed forward and reverse primers at 50uM (good for >6 nL-qPCR chips; \$250)

**Table S2.** Coefficient of variation in target gene copy number across 15-20 chips that were run on two instruments at separate facilities, by two operators at each facility. Synthetic DNA standards were evaluated over a dilution series from 10 to 10^6^ copies/reaction. Coefficient of variation for 10 copies/reaction is not shown when the assay limit of detection was determined to be higher than 10 copies/reaction.

| Organism (target gene) | 10 | 10<sup>2</sup> | 10<sup>3</sup> | 10<sup>4</sup> | 10<sup>5</sup> | 10<sup>6</sup> |
| --- | --- | --- | --- | --- | --- | --- |
| **Pathogenic E. coli** |  |  |  |  |  |  |
| EAEC (aggR) | 115.3 | 40.9 | 26.1 | 25.0 | 16.3 | 61.8 |
| ST-ETEC (STh) |  | 51.1 | 45.0 | 22.7 | 18.4 | 20.5 |
| ST-ETEC (STp) |  | 39.3 | 38.8 | 25.9 | 22.7 | 29.5 |
| LT-ETEC (eltA) |  | 48.7 | 39.4 | 26.8 | 19.8 | 22.2 |
| EPEC (bfpA) |  | 43.3 | 54.2 | 31.8 | 17.2 | 19.8 |
| EPEC (eaeA) | 55.3 | 35.5 | 73.2 | 23.0 | 15.4 | 20.2 |
| STEC (stx1) |  | 47.2 | 18.2 | 21.0 | 13.8 | 16.8 |
| STEC (stx2) |  | 57.3 | 21.8 | 23.5 | 14.6 | 22.8 |
| **Other Bacteria** |  |  |  |  |  |  |
| Campylobacter jejuni/coli (cdtA) |  | 51.6 | 44.4 | 29.7 | 21.8 | 33.3 |
| Clostridium difficile (tcdB) |  | 34.0 | 19.2 | 27.4 | 15.8 | 21.9 |
| Clostridium perfringens (CPE) | 52.1 | 45.8 | 22.4 | 22.8 | 16.0 | 21.7 |
| Helicobacter pylori (ureA) |  | 92.4 | 25.8 | 35.3 | 13.8 | 31.1 |
| Salmonella enterica (invA) | 53.1 | 31.3 | 23.1 | 20.6 | 12.2 | 18.1 |
| Shigella/EIEC (ipaH) |  | 29.4 | 19.4 | 20.4 | 13.8 | 21.2 |
| Vibrio cholerae (tcpA) |  | 44.0 | 22.3 | 25.4 | 15.4 | 18.8 |
| Yersinia enterocolitica (yadA) |  | 60.8 | 101.0 | 29.0 | 16.6 | 21.6 |
| **Protozoa & Helminthes** |  |  |  |  |  |  |
| Cryptosporidium (18S) |  | 102.9 | 112.3 | 40.1 | 31.5 | 19.7 |
| Entamoeba histolytica (18S) |  | 45.9 | 25.6 | 28.6 | 35.6 | 23.7 |
| Giardia (18S) |  | 31.5 | 19.5 | 21.7 | 12.9 | 27.6 |
| Ascaris lumbricoides (ITS1) | 53.2 | 30.6 | 19.1 | 21.6 | 14.7 | 20.2 |
| Trichuris trichiura (18S) | 57.7 | 31.8 | 17.2 | 21.4 | 12.5 | 17.2 |
| **General** |  |  |  |  |  |  |
| Total Archaea (16S) |  | 74.6 | 44.7 | 49.5 | 31.4 | 42.8 |
| Total Bacteria (16S) | 318.9 | 78.2 | 91.0 | 29.7 | 16.7 | 30.5 |
| Total Fungi (ITS1) |  | 54.5 | 40.1 | 53.5 | 44.8 | 40.5 |
| **QC** |  |  |  |  |  |  |
| PhHV (gB) |  | 66.1 | 25.8 | 22.9 | 15.4 | 21.8 |

## References

1. van den Berg RJ, Vaessen N, Endtz HP, Schülin T, van der Vorm ER, Kuijper EJ. 2007. Evaluation of real-time PCR and conventional diagnostic methods for the detection of Clostridium difficile-associated diarrhoea in a prospective multicentre study. J Med Microbiol 56:36–42.

2. Atmar RL, Opekun AR, Gilger MA, Estes MK, Crawford SE, Neill FH, Graham DY. 2008. Norwalk virus shedding after experimental human infection. Emerg Infect Dis 14:1553–7.

3. Liu J, Platts-Mills JA, Juma J, Kabir F, Nkeze J, Okoi C, Operario DJ, Uddin J, Ahmed S, Alonso PL, Antonio M, Becker SM, Blackwelder WC, Breiman RF, Faruque ASG, Fields B, Gratz J, Haque R, Hossain A, Hossain MJ, Jarju S, Qamar F, Iqbal NT, Kwambana B, Mandomando I, McMurry TL, Ochieng C, Ochieng JB, Ochieng M, Onyango C, Panchalingam S, Kalam A, Aziz F, Qureshi S, Ramamurthy T, Roberts JH, Saha D, Sow SO, Stroup SE, Sur D, Tamboura B, Taniuchi M, Tennant SM, Toema D, Wu Y, Zaidi A, Nataro JP, Kotloff KL, Levine MM, Houpt ER. 2016. Use of quantitative molecular diagnostic methods to identify causes of diarrhoea in children: a reanalysis of the GEMS case-control study. Lancet 388:1291–1301.

4. Platts-Mills JA, Liu J, Rogawski ET, Kabir F, Lertsethtakarn P, Siguas M, Khan SS, Praharaj I, Murei A, Nshama R, Mujaga B, Havt A, Maciel IA, McMurry TL, Operario DJ, Taniuchi M, Gratz J, Stroup SE, Roberts JH, Kalam A, Aziz F, Qureshi S, Islam MO, Sakpaisal P, Silapong S, Yori PP, Rajendiran R, Benny B, McGrath M, McCormick BJJ, Seidman JC, Lang D, Gottlieb M, Guerrant RL, Lima AAM, Leite JP, Samie A, Bessong PO, Page N, Bodhidatta L, Mason C, Shrestha S, Kiwelu I, Mduma ER, Iqbal NT, Bhutta ZA, Ahmed T, Haque R, Kang G, Kosek MN, Houpt ER, Acosta AM, Rios de Burga R, Chavez CB, Flores JT, Olotegui MP, Pinedo SR, Trigoso DR, Vasquez AO, Ahmed I, Alam D, Ali A, Rasheed M, Soofi S, Turab A, Yousafzai A, Zaidi AK, Shrestha B, Rayamajhi BB, Strand T, Ammu G, Babji S, Bose A, George AT, Hariraju D, Jennifer MS, John S, Kaki S, Karunakaran P, Koshy B, Lazarus RP, Muliyil J, Ragasudha P, Raghava MV, Raju S, Ramachandran A, Ramadas R, Ramanujam K, Rose A, Roshan R, Sharma SL, Sundaram S, Thomas RJ, Pan WK, Ambikapathi R, Carreon JD, Doan V, Hoest C, Knobler S, Miller MA, Psaki S, Rasmussen Z, Richard SA, Tountas KH, Svensen E, Amour C, Bayyo E, Mvungi R, Pascal J, Yarrot L, Barrett L, Dillingham R, Petri WA, Scharf R, Ahmed AS, Alam MA, Haque U, Hossain MI, Islam M, Mahfuz M, Mondal D, Nahar B, Tofail F, Chandyo RK, Shrestha PS, Shrestha R, Ulak M, Bauck A, Black R, Caulfield L, Checkley W, Lee G, Schulze K, Scott S, Murray-Kolb LE, Ross AC, Schaefer B, Simons S, Pendergast L, Abreu CB, Costa H, Di Moura A, Filho JQ, Leite ÁM, Lima NL, Lima IF, Maciel BL, Medeiros PH, Moraes M, Mota FS, Oriá RB, Quetz J, Soares AM, Mota RM, Patil CL, Mahopo C, Maphula A, Nyathi E. 2018. Use of quantitative molecular diagnostic methods to assess the aetiology, burden, and clinical characteristics of diarrhoea in children in low-resource settings: a reanalysis of the MAL-ED cohort study. Lancet Glob Heal 6:e1309–e1318.

5. Spierings G, Ockhuijsen C, Hofstra H, Tommassen J. 1993. Polymerase chain reaction for the specific detection of Escherichia coli/Shigella. Res Microbiol 144:557–564.

6. Wood MW, Mahon J, Lax AJ. 1994. Development of a probe and PCR primers specific to the virulence plasmid of Salmonella enteritidis. Mol Cell Probes 8:473–479.

7. Lin M-H, Chen T-C, Kuo T-T, Tseng C-C, Tseng C-P. 2000. Real-Time PCR for Quantitative Detection of Toxoplasma gondiiJournal of Clinical Microbiology.

8. He Q, Wang JP, Osato M, Lachman LB. 2002. Real-time quantitative PCR for detection of Helicobacter pylori. J Clin Microbiol 40:3720–3728.

9. Poirier P, Wawrzyniak I, Albert A, El Alaoui H, Delbac F, Livrelli V. 2011. Development and evaluation of a real-time PCR assay for detection and quantification of blastocystis parasites in human stool samples: prospective study of patients with hematological malignancies. J Clin Microbiol 49:975–83.

10. Soumet C, Ermel G, Rose N, Rose V, Drouin P, Salvat G, Colin P. 1999. Evaluation of a multiplex PCR assay for simultaneous identification of Salmonella sp., Salmonella enteritidis and Salmonella typhimurium from environmental swabs of poultry houses. Lett Appl Microbiol 28:113–117.

11. Vet JAM, Majithia AR, Marras SAE, Tyagi S, Dube S, Poiesz BJ, Kramer FR. 1999. Multiplex detection of four pathogenic retroviruses using molecular beacons. Proc Natl Acad Sci 96:6394–6399.

12. Taniuchi M, Verweij JJ, Noor Z, Sobuz SU, Van Lieshout L, Petri WA, Haque R, Houpt ER. 2011. High throughput multiplex PCR and probe-based detection with luminex beads for seven intestinal parasites. Am J Trop Med Hyg 84:332–337.

13. Taniuchi M, Walters CC, Gratz J, Maro A, Kumburu H, Serichantalergs O, Sethabutr O, Bodhidatta L, Kibiki G, Toney DM, Berkeley L, Nataro JP, Houpt ER. 2012. Development of a multiplex polymerase chain reaction assay for diarrheagenic Escherichia coli and Shigella spp. and its evaluation on colonies, culture broths, and stool. Diagn Microbiol Infect Dis 73:121–128.

14. Liu J, Gratz J, Maro A, Kumburu H, Kibiki G, Taniuchi M, Howlader AM, Sobuz SU, Haque R, Talukder K a, Qureshi S, Zaidi A, Haverstick DM, Houpt ER. 2012. Simultaneous detection of six diarrhea-causing bacterial pathogens with an in-house PCR-luminex assay. J Clin Microbiol 50:98–103.

15. Liu J, Gratz J, Amour C, Kibiki G, Becker S, Janaki L, Verweij JJ, Taniuchi M, Sobuz SU, Haque R, Haverstick DM, Houpt ER. 2013. A laboratory-developed TaqMan Array Card for simultaneous detection of 19 enteropathogens. J Clin Microbiol 51:472–80.

16. Liu J, Gratz J, Amour C, Nshama R, Walongo T, Maro A, Mduma E, Platts-Mills J, Boisen N, Nataro J, Haverstick DM, Kabir F, Lertsethtakarn P, Silapong S, Jeamwattanalert P, Bodhidatta L, Mason C, Begum S, Haque R, Praharaj I, Kang G, Houpt ER. 2016. Optimization of quantitative PCR methods for enteropathogen detection. PLoS One 11:1–11.

17. Buss SN, Leber A, Chapin K, Fey PD, Bankowski MJ, Jones MK, Rogatcheva M, Kanack KJ, Bourzac KM. 2015. Multicenter Evaluation of the BioFire FilmArray Gastrointestinal Panel for Etiologic Diagnosis of Infectious Gastroenteritis. J Clin Microbiol 53:915–925.

18. Huang RSP, Johnson CL, Pritchard L, Hepler R, Ton TT, Dunn JJ. 2016. Performance of the Verigene^®^ enteric pathogens test, Biofire FilmArray™ gastrointestinal panel and Luminex xTAG^®^ gastrointestinal pathogen panel for detection of common enteric pathogens. Diagn Microbiol Infect Dis 86:336–339.

19. Wongboot W, Okada K, Chantaroj S, Kamjumphol W, Hamada S. 2018. Simultaneous detection and quantification of 19 diarrhea-related pathogens with a quantitative real-time PCR panel assay. J Microbiol Methods 151:76–82.

20. Platts-Mills J a, Gratz J, Mduma E, Svensen E, Amour C, Liu J, Maro A, Saidi Q, Swai N, Kumburu H, McCormick BJJ, Kibiki G, Houpt ER. 2014. Association between stool enteropathogen quantity and disease in Tanzanian children using TaqMan array cards: a nested case-control study. Am J Trop Med Hyg 90:133–8.

21. J.A. P-M, M. T, Md.J. U, S.U. S, M. M, S.M.A. G, D. M, Md.I. H, M.M. I, A.M.S. A, W.A. P, R. H, E.R. H. 2017. Association between enteropathogens and malnutrition in children aged 6-23 mo in Bangladesh: A case-control study. Am J Clin Nutr 105:1132–1138.

22. Rogawski ET, Liu J, Platts-Mills JA, Kabir F, Lertsethtakarn P, Siguas M, Khan SS, Praharaj I, Murei A, Nshama R, Mujaga B, Havt A, Maciel IA, Operario DJ, Taniuchi M, Gratz J, Stroup SE, Roberts JH, Kalam A, Aziz F, Qureshi S, Islam MO, Sakpaisal P, Silapong S, Yori PP, Rajendiran R, Benny B, McGrath M, Seidman JC, Lang D, Gottlieb M, Guerrant RL, Lima AAM, Leite JP, Samie A, Bessong PO, Page N, Bodhidatta L, Mason C, Shrestha S, Kiwelu I, Mduma ER, Iqbal NT, Bhutta ZA, Ahmed T, Haque R, Kang G, Kosek MN, Houpt ER. 2018. Use of quantitative molecular diagnostic methods to investigate the effect of enteropathogen infections on linear growth in children in low-resource settings: longitudinal analysis of results from the MAL-ED cohort study. Lancet Glob Heal 6:e1319–e1328.

23. Schnee AE, Haque R, Taniuchi M, Uddin MJ, Alam MM, Liu J, Rogawski ET, Kirkpatrick B, Houpt ER, Petri WA, Platts-Mills JA. 2018. Identification of Etiology-Specific Diarrhea Associated with Linear Growth Faltering in Bangladeshi Infants. Am J Epidemiol 187:2210–2218.

24. Grassly NC, Praharaj I, Babji S, Kaliappan SP, Giri S, Venugopal S, Parker EPK, Abraham A, Muliyil J, Doss S, Raman U, Liu J, Peter JV, Paranjape M, Jeyapaul S, Balakumar S, Ravikumar J, Srinivasan R, Bahl S, Iturriza-Gómara M, Uhlig HH, Houpt ER, John J, Kang G. 2016. The effect of azithromycin on the immunogenicity of oral poliovirus vaccine: a double-blind randomised placebo-controlled trial in seronegative Indian infants. Lancet Infect Dis 16:905–914.

25. Taniuchi M, Platts-Mills JA, Begum S, Uddin MJ, Sobuz SU, Liu J, Kirkpatrick BD, Colgate ER, Carmolli MP, Dickson DM, Nayak U, Haque R, Petri WA, Houpt ER. 2016. Impact of enterovirus and other enteric pathogens on oral polio and rotavirus vaccine performance in Bangladeshi infants. Vaccine 34:3068–3075.

26. Stedtfeld RD, Baushke SW, Tourlousse DM, Miller SM, Stedtfeld TM, Gulari E, Tiedje JM, Hashsham S a. 2008. Development and experimental validation of a predictive threshold cycle equation for quantification of virulence and marker genes by high-throughput nanoliter-volume PCR on the OpenArray platform. Appl Environ Microbiol 74:3831–8.

27. Goldfarb DM, Dixon B, Moldovan I, Barrowman N, Mattison K, Zentner C, Baikie M, Bidawid S, Chan F, Slinger R. 2013. Nanolitre real-time PCR detection of bacterial, parasitic, and viral agents from patients with diarrhoea in Nunavut, Canada. Int J Circumpolar Health 72:1–8.

28. Ishii S, Segawa T, Okabe S. 2013. Simultaneous quantification of multiple food-and waterborne pathogens by use of microfluidic quantitative PCR. Appl Environ Microbiol 79:2891–8.

29. Wang F-H, Qiao M, Su J-Q, Chen Z, Zhou X, Zhu Y-G. 2014. High throughput profiling of antibiotic resistance genes in urban park soils with reclaimed water irrigation. Environ Sci Technol 48:9079–9085.

30. Karkman A, Johnson TA, Lyra C, Stedtfeld RD, Tamminen M, Tiedje JM, Virta M. 2016. High-throughput quantification of antibiotic resistance genes from an urban wastewater treatment plant. FEMS Microbiol Ecol 92:1–7.

31. Stedtfeld RD, Williams MR, Fakher U, Johnson TA, Stedtfeld TM, Wang F, Khalife WT, Hughes M, Etchebarne BE, Tiedje JM, Hashsham SA. 2016. Antimicrobial resistance Dashboard application for mapping environmental occurrence and resistant pathogens. FEMS Microbiol Ecol 92:1–9.

32. Mayer-Blackwell K, Azizian MF, Machak C, Vitale E, Carpani G, de Ferra F, Semprini L, Spormann AM. 2014. Nanoliter qPCR Platform for Highly Parallel, Quantitative Assessment of Reductive Dehalogenase Genes and Populations of Dehalogenating Microorganisms in Complex Environments. Environ Sci Technol 48:9659–9667.

33. Kotloff KL, Nataro JP, Blackwelder WC, Nasrin D, Farag TH, Panchalingam S, Wu Y, Sow SO, Sur D, Breiman RF, Faruque AS, Zaidi AK, Saha D, Alonso PL, Tamboura B, Sanogo D, Onwuchekwa U, Manna B, Ramamurthy T, Kanungo S, Ochieng JB, Omore R, Oundo JO, Hossain A, Das SK, Ahmed S, Qureshi S, Quadri F, Adegbola R a, Antonio M, Hossain MJ, Akinsola A, Mandomando I, Nhampossa T, Acácio S, Biswas K, O’Reilly CE, Mintz ED, Berkeley LY, Muhsen K, Sommerfelt H, Robins-Browne RM, Levine MM. 2013. Burden and aetiology of diarrhoeal disease in infants and young children in developing countries (the Global Enteric Multicenter Study, GEMS): a prospective, case-control study. Lancet 6736:1–14.

34. Platts-Mills JA, Babji S, Bodhidatta L, Gratz J, Haque R, Havt A, McCormick BJJ, McGrath M, Olortegui MP, Samie A, Shakoor S, Mondal D, Lima IFN, Hariraju D, Rayamajhi BB, Qureshi S, Kabir F, Yori PP, Mufamadi B, Amour C, Carreon JD, Richard SA, Lang D, Bessong P, Mduma E, Ahmed T, Lima AAAM, Mason CJ, Zaidi AKM, Bhutta ZA, Kosek M, Guerrant RL, Gottlieb M, Miller M, Kang G, Houpt ER, Chavez CB, Trigoso DR, Flores JT, Vasquez AO, Pinedo SR, Acosta AM, Ahmed I, Alam D, Ali A, Rasheed M, Soofi S, Turab A, Yousafzai AK, Bose A, Jennifer MS, John S, Kaki S, Koshy B, Muliyil J, Raghava MV, Ramachandran A, Rose A, Sharma SL, Thomas RJ, Pan W, Ambikapathi R, Charu V, Dabo L, Doan V, Graham J, Hoest C, Knobler S, Mohale A, Nayyar G, Psaki S, Rasmussen Z, Seidman JC, Wang V, Blank R, Tountas KH, Swema BM, Yarrot L, Nshama R, Ahmed AMS, Tofail F, Hossain I, Islam M, Mahfuz M, Chandyo RK, Shrestha PS, Shrestha R, Ulak M, Black R, Caulfield L, Checkley W, Chen P, Lee G, Murray-Kolb LE, Schaefer B, Pendergast L, Abreu C, Costa H, Moura A Di, Filho JQ, Leite ??lvaro, Lima N, Maciel B, Moraes M, Mota F, Ori?? R, Quetz J, Soares A, Patil CL, Mahopo C, Mapula A, Nesamvuni C, Nyathi E, Barrett L, Petri WA, Scharf R, Shrestha B, Shrestha SK, Strand T, Svensen E. 2015. Pathogen-specific burdens of community diarrhoea in developing countries: A multisite birth cohort study (MAL-ED). Lancet Glob Heal 3:564–575.

35. Altschul SF, Gish W, Miller W, Myers EW, Lipman DJ. 1990. Basic local alignment search tool. J Mol Biol 215:403–410.

36. Untergasser A, Cutcutache I, Koressaar T, Ye J, Faircloth BC, Remm M, Rozen SG. 2012. Primer3-new capabilities and interfaces. Nucleic Acids Res 40:e115.

37. Verweij JJ, Blange RA, Templeton K, Schinkel J, Brienen EAT, Rooyen MAA Van, Lieshout L Van, Polderman AM. 2004. Simultaneous Detection of Entamoeba histolytica, Giardia lamblia, and Cryptosporidium parvum in Fecal Samples by Using Multiplex Real-Time PCR. J Clin Microbiol 42:1220–1223.

38. Wiria AE, Prasetyani MA, Hamid F, Wammes LJ, Lell B, Ariawan I, Uh HW, Wibowo H, Djuardi Y, Wahyuni S, Sutanto I, May L, Luty AJF, Verweij JJ, Sartono E, Yazdanbakhsh M, Supali T. 2010. Does treatment of intestinal helminth infections influence malaria? Background and methodology of a longitudinal study of clinical, parasitological and immunological parameters in Nangapanda, Flores, Indonesia (ImmunoSPIN Study). BMC Infect Dis 10:77.

39. Yu Y, Lee C, Kim J, Hwang S. 2005. Group-Specific Primer and Probe Sets to Detect Methanogenic Communities Using Quantitative Real-Time Polymerase Chain Reaction. Biotechnol Bioeng 89:670–679.

40. Hoffmann C, Dollive S, Grunberg S, Chen J, Li H, Wu GD, Lewis JD, Bushman FD. 2013. Archaea and fungi of the human gut microbiome: correlations with diet and bacterial residents. PLoS One 8:e66019.

41. Niesters HG. 2002. Clinical virology in real time. J Clin Virol 25:3–12.

42. Bustin S a, Benes V, Garson J a, Hellemans J, Huggett J, Kubista M, Mueller R, Nolan T, Pfaffl MW, Shipley GL, Vandesompele J, Wittwer CT. 2009. The MIQE guidelines: minimum information for publication of quantitative real-time PCR experiments. Clin Chem 55:611–22.

43. Rutledge RG, Côté C. 2003. Mathematics of quantitative kinetic PCR and the application of standard curves. Nucleic Acids Res 31:e93.

44. Arnold BF, Null C, Luby SP, Unicomb L, Stewart CP, Dewey KG, Ahmed T, Ashraf S, Christensen G, Clasen T, Dentz HN, Fernald LCH, Haque R, Hubbard AE, Kariger P, Leontsini E, Lin A, Njenga SM, Pickering AJ, Ram PK, Tofail F, Winch PJ, Colford JM. 2013. Cluster-randomised controlled trials of individual and combined water, sanitation, hygiene and nutritional interventions in rural Bangladesh and Kenya: the WASH Benefits study design and rationale. BMJ Open 3:e003476.

45. Luby SP, Rahman M, Arnold BF, Unicomb L, Ashraf S, Winch PJ, Stewart CP, Begum F, Hussain F, Benjamin-Chung J, Leontsini E, Naser AM, Parvez SM, Hubbard AE, Lin A, Nizame FA, Jannat K, Ercumen A, Ram PK, Das KK, Abedin J, Clasen TF, Dewey KG, Fernald LC, Null C, Ahmed T, Colford JM. 2018. Effects of water quality, sanitation, handwashing, and nutritional interventions on diarrhoea and child growth in rural Bangladesh: A cluster randomised controlled trial. Lancet Glob Heal 6:30490–4.

46. Stewart CP, Dewey KG, Lin A, Pickering AJ, Byrd KA, Jannat K, Ali S, Rao G, Dentz HN, Kiprotich M, Arnold CD, Arnold BF, Allen LH, Shahab-Ferdows S, Ercumen A, Grembi JA, Naser AM, Rahman M, Unicomb L, Colford JM, Luby SP, Null C. 2018. Effects of lipid-based nutrient supplements and infant and young child feeding counseling with or without improved water, sanitation, and hygiene (WASH) on anemia and micronutrient status: results from 2 cluster-randomized trials in Kenya and Bangladesh. Am J Clin Nutr 109:148–164.

47. Lin A, Ali S, Arnold BF, Rahman MZ, Alauddin M, Grembi J, Mertens AN, Famida SL, Akther S, Hossen MS, Mutsuddi P, Shoab AK, Hussain Z, Rahman M, Unicomb L, Ashraf S, Naser AM, Parvez SM, Ercumen A, Benjamin-Chung J, Haque R, Ahmed T, Hossain MI, Choudhury N, Jannat K, Alauddin ST, Minchala SG, Cekovic R, Hubbard AE, Stewart CP, Dewey KG, Colford JM, Luby SP. 2019. Effects of water, sanitation, handwashing, and nutritional interventions on environmental enteric dysfunction in young children: a cluster-randomized controlled trial in rural Bangladesh. Clin Infect Dis.

48. Schmittgen TD, Livak KJ. 2008. Analyzing real-time PCR data by the comparative CT method. Nat Protoc 3:1101–1108.

49. Hellemans J, Mortier G, De Paepe A, Speleman F, Vandesompele J. 2007. qBase relative quantification framework and software for management and automated analysis of real-time quantitative PCR data. Genome Biol 8:R19.

50. Benjamini Y, Hochberg Y. 1995. Controlling the False Discovery Rate: A Practical and Powerful Approach to Multiple Testing. J R Stat Soc Ser B 57:289–300.

51. Stevenson M with contributions from, Nunes T, Heuer C, Marshall J, Sanchez J, Thornton R, Reiczigel J, Robison-Cox J, Sebastiani P, Solymos P, Yoshida K, Jones G, Pirikahu S, Firestone S, Kyle R, Popp J, Jay M. 2019. epiR: Tools for the Analysis of Epidemiological Data. R Package Version 1.0-2.

52. Lambert P. 2007. Statistical Guidance on Reporting Results from Studies Evaluating Diagnostic Tests.

53. David Collett. 1999. Modelling Binary Data, p. 24. In Modelling Binary Data: Tests in Statistical Science. Chapman & Hall/CRC.

54. Rothman KJ. 2002. No Title, p. 130–143. In Epidemiology: An Introduction. Oxford University Press.

55. Martin Bland J, Altman DG. 1986. Statistical Methods for Assessing Agreement Between Two Methods of Clinical Measurement. Lancet 327:307–310.

56. Datta D. 2017. blandr: a Bland-Altman Method Comparison package for R.

57. Bland DG, Altman JM. 2002. Commentary on Quantifying Agreement between Two Methods of Measurement. Clin Chem 48:801–802.

58. Hasan NA, Ceccarelli D, Grim CJ, Taviani E, Choi J, Sadique A, Alam M, Siddique AK, Sack RB, Huq A, Colwell RR. 2013. Distribution of Virulence Genes in Clinical and Environmental Vibrio cholerae Strains in Bangladesh.

59. Maeda H, Fujimoto C, Haruki Y, Maeda T, Kokeguchi S, Petelin M, Arai H, Tanimoto I, Nishimura F, Takashiba S. 2003. Quantitative real-time PCR using TaqMan and SYBR Green for Actinobacillus actinomycetemcomitans, Porphyromonas gingivalis, Prevotella intermedia, tetQ gene and total bacteria. FEMS Immunol Med Microbiol 39:81–86.

60. Peng X, Nguyen A, Ghosh D. 2018. Quantification of M13 and T7 bacteriophages by TaqMan and SYBR green qPCR. J Virol Methods 252:100–107.

61. Smyth GK, Michaud J, Scott HS. 2005. Use of within-array replicate spots for assessing differential expression in microarray experiments. Bioinformatics 21:2067–2075.

62. Yuan DS, Irizarry RA. 2006. High-resolution spatial normalization for microarrays containing embedded technical replicates. Bioinformatics 22:3054–3060.

63. Wilson IG. 1997. Inhibition and facilitation of nucleic acid amplification. Appl Environ Microbiol 63:3741–3751.

64. Monteiro L, Bonnemaison D, Vekris A, Petry KG, Bonnet J, Vidal R, Cabrita J, Mégraud F. 1997. Complex polysaccharides as PCR inhibitors in feces. J Clin Microbiol 35:995–998.

